# Maximizing Dissimilarity in Resting State detects Heterogeneous Subtypes in Healthy population associated with High Substance-Use and Problems in Antisocial Personality

**DOI:** 10.1101/787606

**Authors:** Rajan Kashyap, Sagarika Bhattacharjee, B.T. Thomas Yeo, SH Annabel Chen

## Abstract

Patterns in resting-state fMRI (rs-fMRI) are widely used to characterize the trait effects of brain function. In this aspect, multiple rs-fMRI scans from single subjects can provide interesting clues about the rs-fMRI patterns, though scan-to-scan variability pose challenges. Therefore, rs-fMRI’s are either concatenated or the functional connectivity is averaged. This leads to loss of information. Here, we use an alternative way to extract the rs-fMRI features that are common across all the scans by applying Common-and-Orthogonal-Basis-Extraction (COBE) technique. To address this, we employed rs-fMRI of 788 subjects from the human connectome project and estimated the common-COBE-component of each subject from the four rs-fMRI runs. Since the common-COBE-component are specific to a subject, the pattern was used to classify the subjects based on the similarity/dissimilarity of the features. The subset of subjects (n=107) with maximal-COBE-Dissimilarity (MCD) was extracted and the remaining subjects (n = 681) formed the COBE-similarity (CS) group. The distribution of weights of the common-COBE-component for the two groups across rs-fMRI networks and subcortical regions was evaluated. We found the weights in the default mode network to be lower in the MCD compared to the CS. We compared the scores of 69 behavioral measures and found 6 behaviors related to the use of marijuana, illicit drugs, alcohol, and tobacco; and including a measure of antisocial personality to differentiate the two groups. Gender differences were also significant. Altogether findings suggested that subtypes exist even in healthy control population and comparison studies (Case vs Control) need to be mindful of it.

## Introduction

The resting state (rs) in which the subjects lie quietly inside the scanner has been widely adopted in neuroimaging research. Research has reliably shown the potential of rs-fMRI data to reveal the intrinsic architecture of the brain (Biswal, Yetkin, Haughton, & Hyde, 1995; Buckner, Krienen, & Yeo, 2013; Fox & Raichle, 2007) and provide important information regarding the brain functions and cognitive abilities (Mennes et al., 2010; Smith et al., 2009; Tavor et al., 2016; Yeo, Tandi, & Chee, 2015). Therefore, researchers are interested to explore how the patterns in rs-fMRI are manifested across individuals with different traits and lifestyle habits. Consequently, there are attempts to link the rs-fMRI to variety of traits and behaviors (Bertolero, Yeo, Bassett, & D’Esposito, 2018; Dubois, Galdi, Han, Paul, & Adolphs, 2018; Finn et al., 2015; Hampson, Driesen, Skudlarski, Gore, & Constable, 2006; Kashyap et al., 2019; Kong et al., 2018; W. Li, Mai, & Liu, 2014; Rosenberg et al., 2016; Smith et al., 2015). In an interesting work by Smith et al., (2015), a positive-negative axis linking various demographic and lifestyle factors to the resting state functional connectivity was delineated. They found positive correlations to positive personal qualities (e.g., high performance on memory and cognitive test, life satisfaction) and negative correlation values for the negative traits (like substance use, anger, rule-breaking behavior).

The way the resting state networks are organized in the brain has been associated to our state, psyche, social values as well as lifestyle habits (Buckner, Andrews-Hanna, & Schacter, 2008; Gusnard & Raichle, 2001; van den Heuvel & Hulshoff Pol, 2010). In this aspect, multiple scans from each subject are helpful and previous research has shown that conclusions from single session could be erroneous (McGonigle et al., 2000) though intersession variabilities exist (McGonigle et al., 2000; Smith et al., 2005). Such intersession variabilities are not only limited to physiological changes in the subjects inside the scanner but also includes alterations in cognitive and affective states (Gorgolewski et al., 2015). With the advent of big rs-fMRI data with multiple scans of each individual, the conventional way either concatenates the rs-fMRI or averages the functional connectivity over the multiple scans. This might lead to loss of important information from the data. An alternate way would be to extract the features of fMRI that are common across all the resting state scans. With this motivation, using the human connectome project (HCP; Smith et al., 2013; Van Essen et al., 2012) data wherein all healthy subjects are scanned four times and performances across numerous behavioral measures are recorded inside and outside the scanner, we aimed to extract the common features shared by the four rs-fMRI runs of a single subject and explore the pattern across all the subjects.

To this aim, we applied the common orthogonal basis extraction (COBE) algorithm (Zhou, Cichocki, Zhang, & Mandic, 2016a; Zhou et al., 2016b); to the four rs-fMRI runs acquired from a single HCP subject. We repeated this for all the subjects (n = 788). The COBE algorithm was originally developed to extract the common and individual features from a “multi-block” data (collection of matrices). The common features are those that are shared by all the blocks whereas the individual features are specific to a block. The number of common components is to be specified by the user. In our case, each run of rs-fMRI of an individual was taken as a block, so application of COBE to the rs-fMRI of all the four runs (multi-block) will decompose the rs-fMRI signals into a linear sum of a number of common COBE component subspace (shared by all runs) and run-specific subspaces (specific to a run). The common component represents the spatial distribution of weights across the brain that are shared by the rs-fMRI’s of an individual across all the scans (Figure 1B). If the features are common to all the resting state scans of an individual, they could in-principle be used to classify the healthy HCP subjects based on the similarity/dissimilarity of the pattern. This is because the physiological parameters in healthy individuals are considered to be in a normal range and a similar expectation from the rs-fMRI features is preconceived. However, this may not be true considering the wide variability in the rs-fMRI features within the healthy individuals. One interesting approach would be to identify a subgroup of healthy HCP subjects whose pattern of common COBE component is different from the group and then find out which behavioral measures account to the differences between the two healthy groups. This will be intriguing given the general trend to classify subjects based on external symptoms (like healthy vs diseased, control vs patient, etc.) does not consider variations within healthy individuals. However, it is undeniable that there can be subtypes within the healthy individuals whose rs-fMRI features might lie on the edges of the spectrum owing to their lifestyle habits, psychological factors, and physiological traits.

**Figure 1.**
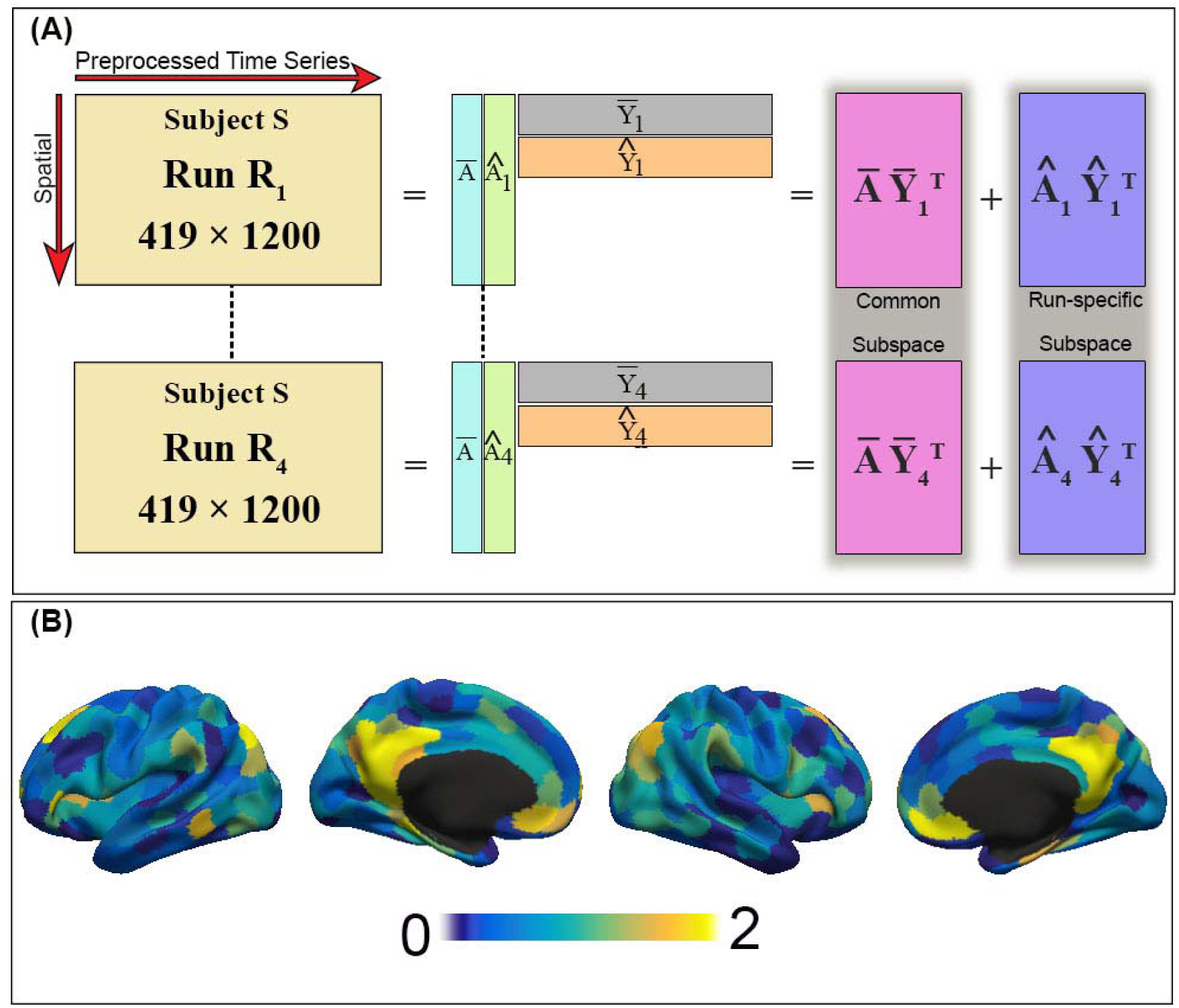
Illustration of Common Orthogonal Basis Extraction (COBE). (A) COBE applied to the 4 runs of rs-fMRI of an HCP subject. COBE projects the rs-fMRI run of an HCP subject onto a common subspace 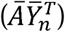 and run-specific subspace 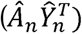. The common component (*Ā*) is shared across all runs. The number of components C spanning the common subspace (i.e. the number of columns of *Ā*) is a user-specified parameter. (B) The spatial map of the weights of the common component *Ā* (419 × C) for C =1 across the cortical areas is shown for a random HCP subject.

Conceptually, it is worth mentioning our work along with the vast literature that had investigated trait-level and state-level aspects of fMRI (Bijsterbosch et al., 2017; Cole, Bassett, Power, Braver, & Petersen, 2014; Gratton et al., 2018; Kong et al., 2018; Krienen, Yeo, & Buckner, 2014; Shirer, Ryali, Rykhlevskaia, Menon, & Greicius, 2012; Wang, Ong, Patanaik, Zhou, & Chee, 2016; Yeo, Krienen, et al., 2015). In our previous work (Kashyap et al., 2019), COBE was applied to the rs-fMRI of each run in the HCP data. Each run comprised of rs-fMRI from 803 subjects. Interestingly, the common COBE component of the four runs revealed the state-effects specific to that run. Subsequently, removing the state-effects from the rs-fMRI improved the functional connectivity based behavioral prediction. Here, our goal was to obtain the common COBE component for all the HCP subjects and classify those subjects whose common COBE components were as dissimilar as possible. Subsequently, we explored the traits that are different in that subset of subjects.

## Methods

### Overview

The rs-fMRI of 1094 subjects from the Human Connectome Project (HCP) were considered in the study. However, only 788 subjects survived the selection criteria and were preprocessed. Thus, COBE was applied to the four rs-fMRI scans of these subjects. The first common component of COBE was extracted for every subject. The common components were correlated to form the common-correlation matrix. The subjects with a dissimilar pattern were then extracted from the matrix and grouped together to form the Maximal-COBE-Dissimilarity (MCD) group. The MCD group and the group of remaining subjects were evaluated for differences in the scores of 62 behavioral measures and 7 physiological parameters.

### Rs-fMRI data

The HCP S1200 release has a multi-modal collection of data across behavioral, structural MRI, resting-state and task-state fMRI, and MEG paradigms from healthy adults (Smith et al., 2009; Van Essen et al., 2012). All imaging data were collected on a custom-made Siemens 3T Skyra scanner that uses a multiband sequence. The rs-fMRI was acquired in 2mm isotropic resolution with a TR of 0.72 seconds for a total of 1200 frames lasting 14 minutes and 33 seconds. The rs-fMRI and behavioral data were collected on two sessions. Each rs-fMRI session consisted of two runs. Therefore, in total each subject had 4 runs of resting state scans.

### Preprocessing

The dataset used in the present study was taken from our previous work (Kashyap et al., 2019). The dataset comprised of the MSMAll ICA-FIX data on fs_LR32K surface space (HCP S1200 manual; Glasser et al., 2013; Griffanti et al., 2014; Salimi-Khorshidi et al., 2014) from 1094 subjects. As few studies have pointed out that ICA-FIX cannot completely remove the head-motion artifacts, so further nuisance regression is necessary (Burgess et al., 2016; Kong et al., 2018; Li et al., 2019; Siegel et al., 2017). Therefore an estimate of framewise displacement (FD; Jenkinson, Bannister, Brady, & Smith, 2002) and root-mean-square of voxel-wise differentiated signal (DVARS) (Power, Barnes, Snyder, Schlaggar, & Petersen, 2012) was done using fsl_motion_outliers. Consequently, those volumes for which FD > 0.2mm and DVARS > 75, as well as uncensored segments of data lasting fewer than 5 contiguous volumes were flagged as outliers. Nuisance regression was performed with 18 regressors that consisted of a global signal, six motion parameters, averaged ventricular signal, averaged white matter signal, and their temporal derivatives. While computing the regression coefficients in nuisance regression, outlier volumes were ignored. Finally, a bandpass filter (0.009 Hz ≤ f ≤ 0.08 Hz) was applied to the data and the BOLD runs with more than half the volumes flagged as outliers were completely removed. As a result, 82 subjects had all runs removed and were thus eliminated.

Adopting the 400 cortical parcellation by Schaefer et al., (2018), we obtained the preprocessed fMRI time courses from each of the 400 cortical parcels. Additionally, 19 subcortical regions (brain stem, accumbens, amygdala, caudate, cerebellum, diencephalon, hippocampus, pallidum, putamen, and thalamus; Fischl et al., 2002) were included. Thus, there were 419 regions in total, and therefore the rs-fMRI of each subject in each run was a 419 × 1200 matrix.

### Behavioral data

We considered a set of 62 behavioral measures and 7 physiological parameters spanning across 8 categories including-alertness, cognition, emotion, personality, sensory, health and family history, psychiatric and life function, and substance use (Table S1). These categories provided a considerable range of the participant’s traits, lifestyle habits, age, body mass index, and health. Importantly, our analyses were restricted to the subjects who had all four runs surviving the quality control procedure with all 62 behavioral measures and 7 physiological parameters, resulting in a final set of 788 subjects.

### Extraction of Common COBE Component

For more details about the COBE algorithm, we refer readers to methodological papers (Zhou et al., 2016a; Zhou et al., 2016b). Here we briefly describe how COBE was applied to rs-fMRI data in this study.

We applied COBE to the rs-fMRI’s of each subject separately. For ease of explanation, let us consider the rs-fMRI (419 × 1200 matrix) of each run as a block. Here, we would like to recall that our study included only those subjects that had 4 runs of resting state scan. For a subject S, let *R*_*n*_ denote the 419 × 1200 rs-fMRI matrix for the n^th^ run (where n is a positive integer with n ≤ 4). As illustrated in Figure 1A, COBE seeks to decompose *R*_*n*_ into

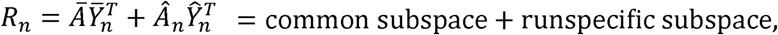

where *Ā* is a 419 × C matrix representing the common component shared across all runs. C is the number of common components, and is defined a priori by the user. In the present study, we evaluated the common COBE component *Ā* of all the subjects (n = 788) for C = 1. The common COBE component for C = 2 and 3 were also evaluated but not considered further in the analysis (see results). A useful property of COBE is that if COBE was applied twice (sequentially) with C = 1, the two common components will be (in practice) the same as the common components obtained by applying COBE once with C = 2 (Zhou et al., 2016a). This has also been tested in our previous work (Kashyap et al, 2019). We mention this property here since one might assume that a minor difference in the data might affect the decomposition and the components may split up differently; which however is not the case with COBE.

The spatial map of the common COBE component for a subject can be visualized in Figure 1B. The 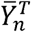 (C × 1200 matrix) is the time course associated with the component. 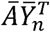 is a 419 × 1200 matrix is representing the common subspace shared by all the runs, while 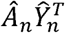 is also a 419 × 1200 matrix representing the projection of the -th run rs-fMRI onto the run-specific subspace.

### Common Correlation matrix

The common component ***Ā*** for every subject (n = 788) was extracted. To classify subjects based on their similarity/dissimilarity of the pattern of the common component, we correlated the component across the subjects. Thus, a 788 × 788 common-correlation matrix was computed using Pearson’s correlation. We seek to select a subset of subjects from the correlation matrix whose common components are as different as possible (i.e. the subject should have a COBE pattern dissimilar to other subjects). To this end, we applied the strategy developed by (Kong et al., 2018). Here, we randomly picked an entry in the common-correlation matrix (i.e., pair of subjects) whose absolute correlation was less than a threshold of 0.75 (also see supplement for the other thresholds that we tested to ascertain 0.75 as our choice for optimal threshold). We continued adding new random subjects, such that each newly added subject was minimally correlated (absolute r < 0.75) with the current set of subjects. The procedure terminated when no more subject could be added. The procedure was repeated 1000 times, resulting in 1000 sets with varied number of subjects per set. Of these 1000 sets, we chose the set containing subjects with the smallest maximum absolute correlation. We will refer to this subset of subjects as the Maximal-COBE-Dissimilarity (MCD) group. Subsequently, the remaining subjects formed the COBE-Similarity (CS) group.

### Behavioral differences between groups

Here we would like to remind the readers that the HCP dataset which is from a healthy population is been dissociated to extract a subset of subjects with dissimilar rs-fMRI COBE pattern. The purpose is to characterize the traits associated with the dissimilarity. For this, we compared the two groups (MCD and CS) across 62 behavioral measures and 7 physiological parameters from 8 categories namely health, alertness, cognition, emotion, personality, psychiatry and life function, sensory, and substance use. The behavioral measures included in each category provided comprehensive information on an individual’s physical health, psyche, and lifestyle habits. For example, the *personality* includes behavioral measure like-agreeableness, open to experiences; *emotion* includes happiness, anger, fear; *substance use* include his/her intake of alcohol, tobacco, illegal drugs, marijuana; *sensory* encompasses the sense of odor, taste, pain; *cognition* involves scores of mini-mental state examination, sleep, working memory, fluid intelligence; and *psychiatric function* include the scores of antisocial personality problem, externalizing, internalizing; and *physiological parameters* include a person’s blood pressure, body mass index, etc. (Table S1).

### Code availability

The code of COBE can be downloaded at http://www.bsp.brain.riken.jp/~zhougx/cifa.html. The code for elastic-net is available freely at https://web.stanford.edu/~hastie/glmnet_matlab/. The Schaefer’s 400-region parcellation is available at https://github.com/ThomasYeoLab/CBIG/tree/master/stable_projects/brain_parcellation/Schaefer2018_LocalGlobal.

## Results

### Overview

COBE was applied to rs-fMRI of 788 HCP subjects to obtain their first common component. The common COBE component of each subject was correlated with the common component of other subjects to obtain the 788 × 788 common-correlation matrix. From this matrix, a subset of subjects that had a dissimilar pattern of common COBE component was extracted to form the MCD group (n = 107) and remaining subjects formed the CS group (n = 681) group. The spatial map of the common component from the MCD and CS group were evaluated and significant differences were found in default mode network. Finally, we also explored the scores of 62 behavioral measures and 7 physiological parameters between the two groups and found only 6 behavioral measures to differentiate the two groups.

### Spatial maps of common COBE components

Common COBE components are illustrated in a spatial map in Figure 2 below. Figures 2A and 2B show the 400 cortical and 19 subcortical ROIs used to compute the 419 × 1200 rs-fMRI time courses. Parcel colors correspond to 17 large-scale-networks (Yeo et al., 2011). The 17 networks are a finer division of the 7 cortical networks (Temporo Parietal, Default mode, Control, Limbic, Attention, Somato-motor, and Visual). The distribution of the weights of the common component (419 × C) for C =1 (i.e. the first common component) for all the subjects across the 7 cortical networks (Yeo et al., 2011) of both hemispheres and subcortical areas can be visualized in Figure 2C. In line with previous studies, we find that the pattern with higher weights in the default mode network is consistent across the subjects. However, visual inspection also reveals that in a few subjects the distribution of weights was different. Therefore, we extracted the subjects with a dissimilar pattern of the common COBE component from the common correlation matrix.

**Figure 2.**
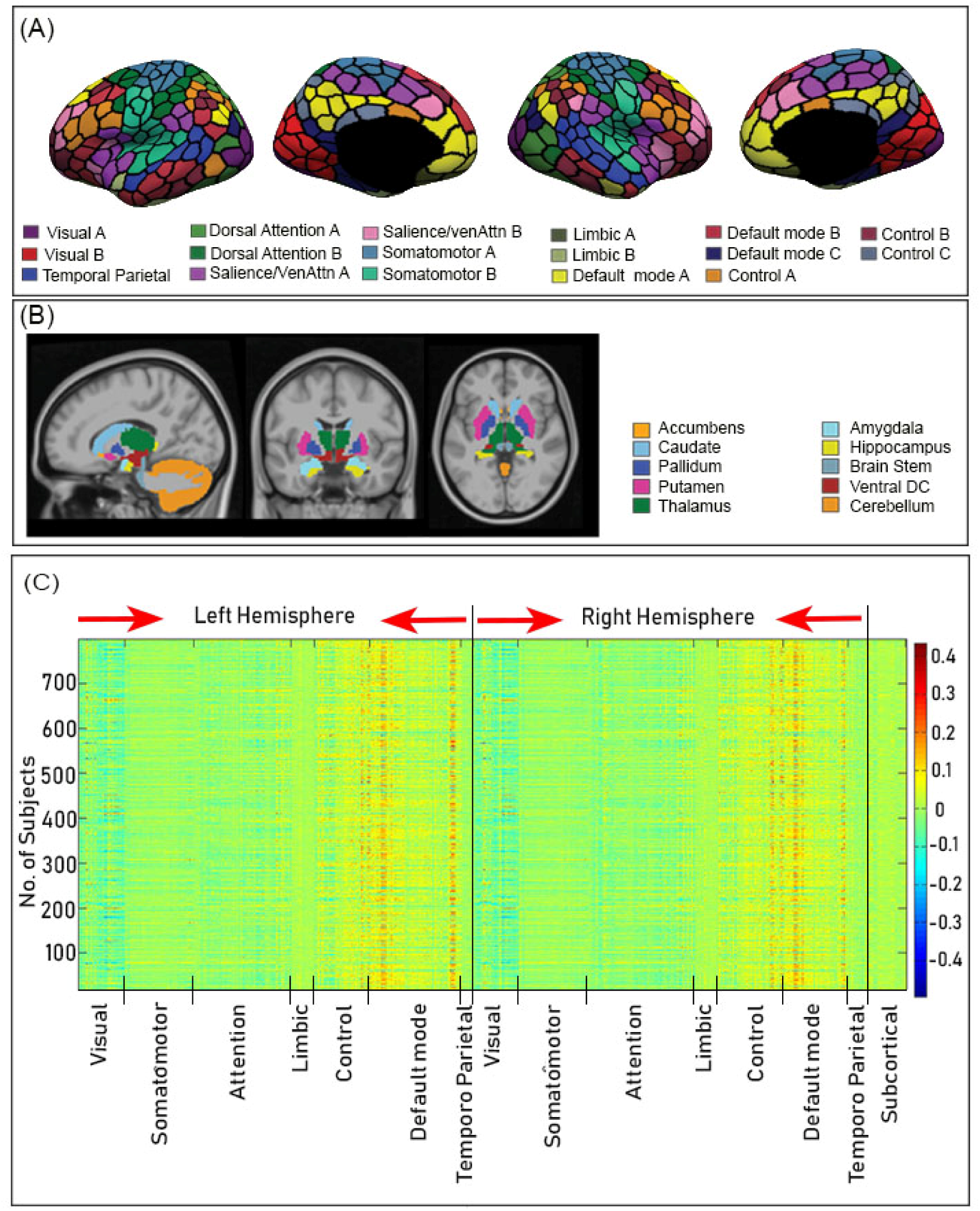
Spatial distribution of common COBE component (C = 1) across the cortical and subcortical areas. (A) 400 cortical parcels (Schaefer et al., 2018). Parcel colors correspond to 17 large-scale-networks (Yeo et al., 2011). The 17 networks are a subdivision of seven networks (Temporal Parietal, Default mode, Control, Limbic, Attention, Somato-motor, and Visual). (B) 19 subcortical ROIs (Fischl et al., 2002). (C) Spatial distribution of weights of the first common COBE component of 788 subjects across the 7 networks (in both hemispheres) and 19 subcortical areas. Most subjects show higher weights in the default mode network areas, though in few subjects the distribution of weight is different.

Interestingly, we also observed the distribution of weights across all the subjects for the second and third common COBE component, i.e. for C = 2 and 3 respectively (supplement). We found that as the number of components increases the similarity of the pattern across the subjects decrease. Visual inspection reveals high inter-individual variation in the distribution of the 2^nd^ and 3^rd^ common COBE component across the 7 resting state networks. Overall, the results suggested not to use higher components to classify the subjects.

The MCD group comprised of 107 subjects (max |r| = .74). Consequently, the CS group had 681 subjects (max |r| = .98). The spatial map of the weights of common COBE component for the two groups (averaged across the subjects) can be visualized in Figure 3A. A clear difference between the two groups can be visualized in the default mode network (DMN) areas especially in the posterior cingulate cortex (PCC). Since, Yeo et al, (2011) have subdivided the DMN into DMN-A, DMN-B, and DMN-C (Figure 2A); so we narrowed down our analysis to the subnetworks of DMN and averaged the weights of the common component across the subjects in the two groups for the 17 cortical networks and 19 subcortical areas (Figure 3B). Specifically, we found that the two groups differed in their distribution of the weights only across DMN-A areas (p < 1*e* ^−14^).

**Figure 3.**
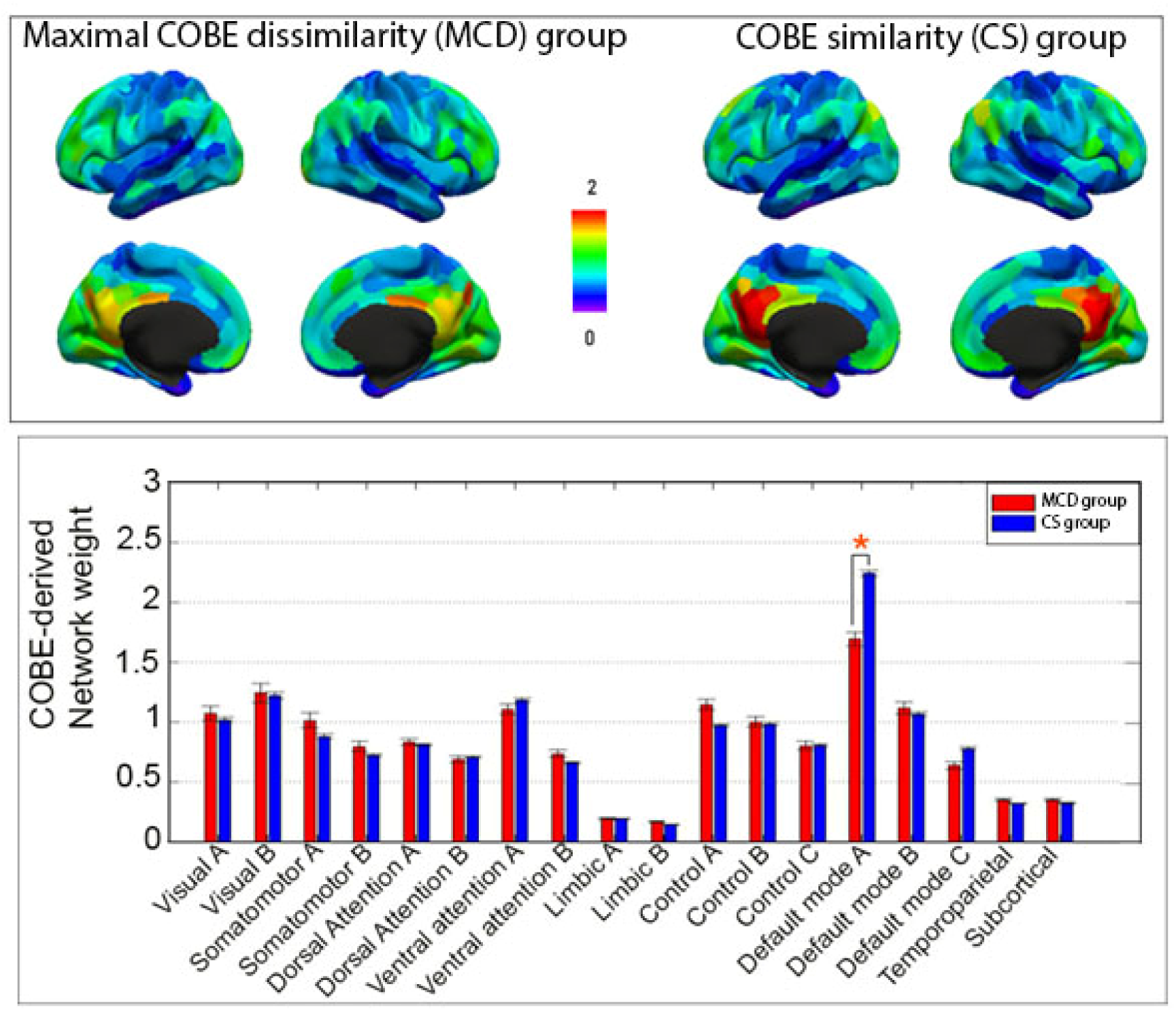
Differences in weight of common component across the default mode network areas between the MCD and CS group. (A) Higher weights of common COBE component can be observed in the Posterior cingulate cortex for the CS group (B) The averaged weights of common COBE component across the 17 cortical networks (Yeo et al., 2011) and subcortical areas is shown for the two groups. * represents the network that survived Bonferroni correction (p < 1*e* ^−14^). Clearly, the MCD group has reduced weights in the DMN-A.

### Behavioral differences between the groups

Scores of 62 behavioral measures and 7 physiological parameters were compared between the MCD and CS group (Table S1). We found 5 behavioral measures to survive the significance level with Bonferroni correction (p < 1*e*^−5^). Interestingly, these behaviors belonged to the two categories (i) *substance use*, and (ii) *psychiatric and life function*. In the first category, we found the behavioral measure that reported a subject’s alcohol consumption in past 7 days (Total_Drinks_7_Days) to be higher for MCD (10.11 ± 1.00) than the CS group (4.84 ± 0.26). Similarly, the tobacco consumption in the past 7 days (Total_Any_Tobacco_7_Days) for MCD (18.05 ± 3.26) was also higher than the CS (5.97 ± 0.76). For the ease of representation, we divided the intake of alcohol and tobacco into five categories based on the number of times they (alcohol/tobacco) were reported to have been consumed by a subject in the past 7 days. They are – 0 (no intake of alcohol/tobacco), 1-5, 6-10, 11-15, 16-20, and > 20 times (Figure 4A and 4B). 39% of subjects in MCD reported ≥ 6 drinks of alcohol/week; which is only 10.7% in the CS. Similarly, 87% in CS group reported to have never taken any kind of tobacco (for e.g., cigarettes, cigars, pipes); which in contrast is only 49.5% in MCD.

**Figure 4.**
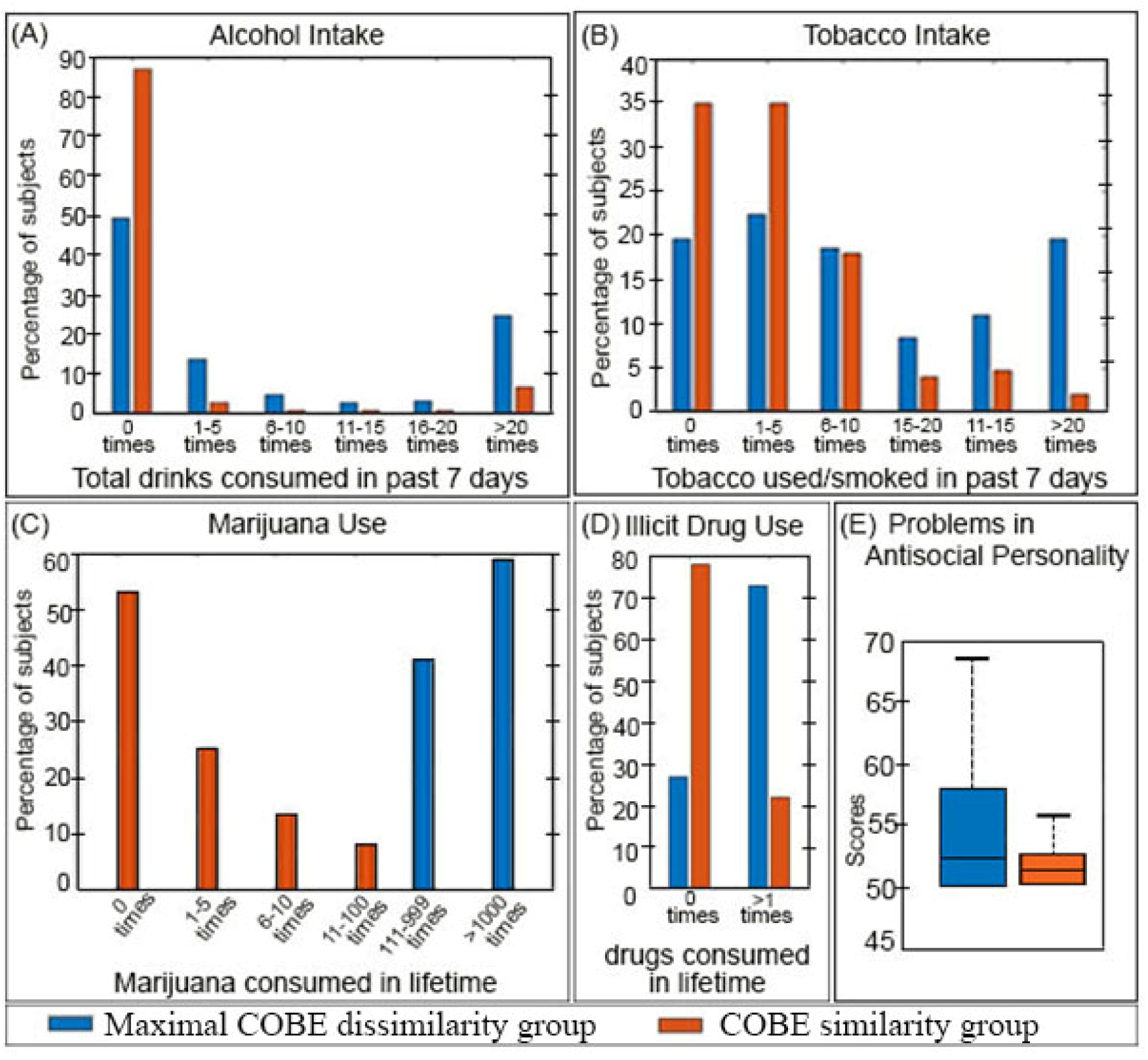
Comparison of behavioral measures between MCD and CS group for the use of-(A) Alcohol (B) Tobacco (C) Marijuana and (D) Illicit drugs; and for (E) Antisocial personality problems.

From the first category, we also found the MCD group to be comprised of those subjects who had a high use of Marijuana in their lifetime (4.59 ± 0.04; SSAGA_Mj_Times_Used) compared to the CS group (1.35 ± 0.06). The HCP has divided the use of Marijuana into five categories based on the number of times it was reported to have been used by the subjects in their lifetime. They are 0 (never used), 1-5, 6-10, 11-100, 111-999, > 1000 times. As shown in Figure 4C, all the subjects in MCD reported having used marijuana for more than 100 times; of which, 59% reported a marijuana intake of more than 1000 times. In contrast, 53% of subjects in the CS group reported to have never used Marijuana, and the majority of the remaining subjects (45%) reported to have used it for less than 100 times. Similarly, the use of illicit drugs (cocaine, hallucinogen, opiates, sedatives, and stimulants; SSAGA_Times_Used_Illicits) was less in CS (0.53 ± 0.04) compared to MCD (2.19 ± 0.16). 78% of the subjects in the CS group reported having never used any illicit drugs. This was only 27% in the MCD (Figure 4D).

In the psychiatric and life function category of behaviors, the score of antisocial personality problems measured by DSM criteria (DSM_Antis_Pct) was marginally higher in MCD than in the CS (Figure 4E). The antisocial personality in HCP is Achenbach Adult Self Report (ASR) questionnaires for the DSM-Oriented Scale that measures a person’s aggressiveness, law-abiding behaviors, etc. Interestingly, since sex related differences are reported for both substance use (Cooper & Craft, 2018) and antisocial personality problems (Cale & Lilienfeld, 2002), we calculated the number of males and females in each group. MCD group comprised of 74 males and 33 females whereas CS group comprised of 323 males and 358 females, respectively. Chi square test revealed gender differences between the groups to be significant (p < 1*e* ^−5^).

Finally, as an additional analysis we also correlated the variation of the common COBE component of each subject in the MCD group with five behavioral measures-Marijuana intake, Illicit Drug use, Alcohol and Tobacco use, and Antisocial personality problems (see supplement).

## Discussion

In this paper, we utilized an alternate way to analyze the multiple scans of rs-fMRI data from an individual. We applied the COBE technique (Zhou et al., 2016a; Zhou et al., 2016b) to decompose the four runs of rs-fMRI of each HCP subject into a common component space (C = 1) shared by the runs of an individual and run-specific subspaces. We observed the distribution of the weights of the first common COBE component across the cortical and subcortical areas for all the HCP subjects (n= 788). Consistent with previous rs-fMRI studies, we found higher weights across the areas of DMN. By visual inspection, distribution of weights across the areas in few subjects appeared dissimilar. Therefore, we attempted to extract the subset of subjects whose common components are as different as possible (Kong et al., 2018). This subset of subjects (n =107) comprised the MCD, and the remaining subjects (n = 681) formed the CS group. We evaluated the distribution of weights of the common COBE component for the two groups across the 17 resting state networks (Yeo et al., 2011) and 19 subcortical areas. We found the weights in the DMN-A to be lower (p < 1*e* ^−14^) in MCD compared to the CS group (Figure 2E). To our knowledge, none of the studies have applied COBE to multiple runs of rs-fMRI of healthy individuals and classified them in subgroups. It is important to mention that appropriate feature extraction is a prerequisite to pattern recognition and classification. While few researchers have used similar techniques like canonical correlation analysis (CCA) to extract features that were hierarchically clustered to identify subtypes of depression (Drysdale et al, 2017), and to delineate positive-negative axis linking various demographic and lifestyle factors (Smith et al, 2015); others have used alternate methods like partial least square (PLS; Krishnan, Williams, McIntosh, Abdi, 2011) and joint and individual variation explained (JIVE; Yu, Risk, Zhang, & Marron, 2017) to establish the relationship between brain activity and behavioral measure. Though JIVE has been suggested as an improved alternative to PLS and CCA (for details, Yu et al, 2017), COBE was shown to be better in estimating the common components compared to JIVE (for details, Zhou et al, 2016a).

Altogether our COBE based approach highlighted that there exists heterogeneity between participants even in healthy control populations. Such heterogeneities are often ignored in comparison studies (healthy vs disease, control vs patient, etc.) because the behavioral manifestations of the observed heterogeneities are not well mapped. To this, we looked for the traits that are different between the two groups. We compared the two groups for 62 behavioral measures and 7 physiological parameters spanning 8 categories that include an individual’s lifestyle habits, cognitive skills, and physical health. Four behavioral measures (use of -marijuana, -illicit drugs, -alcohol, and -tobacco) from the *substance use* category, and one behavioral measure (antisocial personality problem) from the *psychiatric and life function* category had significant difference (p < 1*e* ^−5^) between the two groups. The findings suggested that high substance use and problems of antisocial personality might alter the pattern of common COBE component shared by multiple scans of rs-fMRI. However, it is equally plausible that the observed alterations in common COBE component might indicate vulnerability towards substance use and antisocial personality problems. Interestingly, we also found sex differences between the two groups, with percentage of males being higher in the MCD. Never the less, the existence of subtypes within healthy subjects highlighted that the general categorization of subjects based on external symptoms (e.g. healthy vs diseased, control vs patient etc.) might need to consider aspects of a subject’s lifestyle habits, and psychological factors as they have an impact on the rs-fMRI features.

It was interesting to find that MCD group reported higher use of Marijuana, illicit drugs, alcohol, and tobacco. This was coherent with our observation of reduced weight of the common COBE components in the PCC constituting the core region of DMN-A. Task-based fMRI meta-analysis has reported PCC areas in the DMN to be affected in substance seekers (Blest-Hopley, Giampietro, & Bhattacharyya, 2018; Yanes et al., 2018). The rs-fMRI analysis revealed that longer use of cannabis is associated with decreased low frequency power of the DMN (Thijssen et al., 2017). The posterior cingulate cortex (PCC), a key region in the DMN is responsible for self-consciousness and self-referential mental thoughts (Andrews-Hanna, Reidler, Sepulcre, Poulin, & Buckner, 2010; Andrews-Hanna, Smallwood, & Spreng, 2014; Buckner & Carroll, 2007; Cavanna & Trimble, 2006; Fransson & Marrelec, 2008; Leech & Sharp, 2013; Raichle et al., 2001; Spreng, Mar, & Kim, 2008). Marijuana users were also observed to have alteration in the PCC (Kim et al., 2011). Similarly, individuals with alcohol use disorder, (Chanraud, Pitel, Pfefferbaum, & Sullivan, 2011) reported desynchronized low frequency rs-fMRI signals from the PCC and cerebellum. Nicotine (a key ingredient of tobacco) was also associated with decreased activity in the regions of DMN (Tanabe et al, 2011) and low overall functional connectivity (Cheng et al., 2019). From the vast pool of scientific studies, it can be inferred that tobacco and drugs intake, along with marijuana and alcohol use are associated with differences in resting state time courses (Cheng et al., 2014), and are possibly making adolescents more vulnerable to psychiatric and other disorders (Thijssen et al., 2017).

The subjects in the MCD were found to have a predisposition towards antisocial personality problems. It is important to mention here that the interrelationship between substance use, tobacco-intake, alcoholism, and antisocial behavior is complex (Helstrom, Bryan, Hutchison, Riggs, & Blechman, 2004). These factors are highly correlated both cross-sectionally and across the lifespan, share common risk factors, and are predictive of negative life outcomes (Compton, Conway, Stinson, Colliver, & Grant, 2005; Kendler, Prescott, Myers, & Neale, 2003; Krueger, Markon, Patrick, Benning, & Kramer, 2007; Moffitt, 2017; Windle, 1990). About 40% to 50% of individuals with a substance use disorder meet the criteria for antisocial personality disorder and approximately 90% of individuals diagnosed with an antisocial personality disorder also have a co□occurring substance use disorder (Messina, Farabee, & Rawson, 2003). Results have also suggested that smoking and alcohol use act as mediators between externalizing problems and marijuana and other drug use (Helstrom et al., 2004). Adolescents with antisocial behavior have reported reduced connectivity in the DMN (Broulidakis et al., 2016; Zhou, Yao, et al., 2016). This is not surprising because DMN has an indispensable role in social understanding of others (Li et al., 2014) and the areas responsible for social cognition partly overlap with the DMN (Corbetta, Patel, & Shulman, 2008; Mars et al., 2012; Schilbach, Eickhoff, Rotarska-Jagiela, Fink, & Vogeley, 2008).

Even though we identified the common component shared by multiple runs of rs-fMRI of an individual and observed its distribution across all networks; it is difficult to conclude whether the lower weights in the DMN-A areas accounted for the observed behavioral differences. In a longitudinal study from a large sample (n = 1293) of subjects, it was found that adolescents who reported more antisocial behavior at age 14 were more likely to smoke daily, to drink heavily, and to use illicit drugs at age 17 (Adalbjarnardottir & Rafnsson, 2002). Valuing the association between antisocial personality and other behaviors reported in existing literature, we investigated the correlation between the scores of antisocial personality with other 4 behavioral measures and found weak correlation for all the pairs (see supplement). Never the less, considering that such causal relationship between antisocial behavior and addiction is strong, future studies could investigate whether preexisting differences in DMN leads to antisocial behavior in individuals or whether the differences in DMN are the effect of antisocial behavior. With that said, the present study was an exploratory analysis that found meaningful differences in rs-fMRI features within subgroups of healthy individuals that had the distinctions translated to behavioral measures. The groups differed in their ratio of number of males to females, clearly reflecting that the MCD group comprised more males (see Table S1). This also reflected that the rate of substance and antisocial personality problems are higher in males. This is in accordance with results reported recently using HCP data (Petker et al, 2019), and previous datasets (Cooper & Craft, 2018; Cale & Lilienfeld, 2002). Most importantly, these findings highlight the importance of considering between-subject heterogeneity even in healthy control populations (Kebets et al, 2019); and conveys a subtle message to the comparison studies (healthy vs disease, control vs patient, etc.) to recruit healthy/control subjects based not only on the features relevant to their study, but to also consider the factors that might lead to disparateness in subject’s habit and psyche.

Here we would also like to point out few limitations of the study. From the data perspective, it would have been interesting to obtain additional information about the last use of substance, marijuana as well as tobacco and alcohol. We mention this because an individual’s substance use status (dose, duration etc.) at the time of scanning can affect the COBE indices. In the absence of such vital information it is difficult to rationalize the state versus trait effects of substance use on the common COBE component of a subject. From the approach perspective, we would like to point out that only cortex based parcellation was used in our analysis. Subcortical structures as well as cerebellum were considered as 19 individual nodes in the analysis. We cannot deny the crucial role these vital structures have in substance use and personality disorders (Manza, Tomasi, & Volkow, 2018). We would therefore advise future studies to use integrated parcellations that detail the cortex, cerebellum and the subcortical structures. Keeping these limitations in mind, we look forward to applying the present method on patient groups to identify meaningful disorder subtypes (Zhang et al, 2016; Drysdale et al, 2017).

Finally, it is important to note that the results obtained by our present study are consistent with the results obtained by Smith et al, (2015), though they adopted a different methodology. They applied CCA on the resting state functional connectivity (RSFC) matrices of 461 HCP subjects with 158 behavioral measures to obtain a single mode of brain-behavior covariation. They found that vast majority of the positive behaviors (for e.g., education, income, IQ, life-satisfaction, etc.) correlate positively with this mode, and similarly sets of negative behaviors (for e.g., anger, sleep quality, alcohol intake, substance abuse, Marijuana use, etc.) correlate negatively. Interestingly, they also found DMN regions mainly the medial frontal and parietal cortex in the temporo-parietal junction and in anterior insula and frontal operculum to contribute strongly. It cannot be denied that the CCA mode provided a distinct overview of the brain-behavior organization, however the ability to generate explainable insights at the individual level (subject or behavior specific) was limited. For example, the set of areas belonging to DMN were found to be involved across a wide range of behaviors that surfaced from the first CCA mode; however association of certain behaviors (reading, aggression, working memory, etc.) with these exclusive areas might be of debate. In contrast, our study had a tapered point of view. Using the common signal from the multiple fMRI runs, we focused to delineate subgroups within healthy individuals whose resting state pattern (mainly in the PCC area of DMN) was dissimilar. The dissimilarity translated to significant differences across 6 (out of 69) negative behaviors (including gender), which were also a part of the behavioral measures found significant in the CCA mode (Smith et al, 2015). It is noteworthy to mention that two exploratory studies reporting similar resting-state fMRI-behavior relationship by using different methodologies emphasize the reliability of the findings. Last but not least, the reproducibility of the prime findings is also one of the important purposes of generating big public datasets.

## Conclusions

In this work, we decomposed the rs-fMRI runs of a single HCP subject to extract the common component shared by their multiple runs. We classified those subjects whose common component are as different as possible. In those subjects, we found that the strength of the common component was reduced in the default mode network areas. Interestingly, we also found these subjects to have higher usage of marijuana, illicit drugs, alcohol, and tobacco, and have a predisposition towards antisocial personality problems. The existence of subtypes within healthy individuals that have meaningful differences in their resting-state patterns conveys that the general categorization of subjects based only on external symptoms (e.g. healthy vs diseased, control vs patient, etc.) should also consider aspects of a healthy subject’s lifestyle habits and psyche. Overall the exploratory approach and the findings convey the importance of considering the factors associated with between-subject heterogeneity in healthy control populations.

## Acknowledgment

This work was supported by Nanyang Technological University start-up grant (NTU-SUG). The work was also funded by NUS Strategic Research (DPRT/944/09/14), NUS SOM Aspiration Fund (R185000271720), Singapore NMRC (CBRG/0088/2015), NUS YIA and the Singapore National Research Foundation (NRF) Fellowship (Class of 2017). Our research also utilized resources provided by the Center for Functional Neuroimaging Technologies, P41EB015896 and instruments supported by 1S10RR023401, 1S10RR019307, and 1S10RR023043 from the Athinoula A. Martinos Center for Biomedical Imaging at the Massachusetts General Hospital. Our computational work was partially performed on resources of the National Supercomputing Centre, Singapore (https://www.nscc.sg). Data were provided by the Human Connectome Project, WU-Minn Consortium (Principal Investigators: David Van Essen and Kamil Ugurbil; 1U54MH091657) funded by the 16 NIH Institutes and Centers that support the NIH Blueprint for Neuroscience Research; and by the McDonnell Center for Systems Neuroscience at Washington University.

