## supplement for "Maximizing Dissimilarity in Resting State detects Heterogeneous Subtypes in Healthy population associated with High Substance-Use and Problems in Antisocial Personality"

**Supplemental Results**

Rajan Kashyap_,_ Sagarika Bhattacharjee, B.T.Thomas Yeo, SH Annabel Chen.

*Spatial distribution of Common COBE Components*

The spatial distribution of the weights of the second and third common component of COBE across the 7 cortical networks and 19 subcortical regions have been shown in Figure S1 (A and B), respectively. Visual inspection of the pattern of distribution across all the subjects for both 2^nd^ and 3^rd^ component reveals high inter-individual variation across the networks compared to the 1^st^ common COBE component (Figure 2C). So, classifying subjects whose common COBE component is as different as possible based on the pattern of distribution of 2^nd^ and 3^rd^ common COBE component will be infeasible. Moreover, obtaining an optimal threshold value for the 2^nd^ and 3^rd^ components might also be difficult.

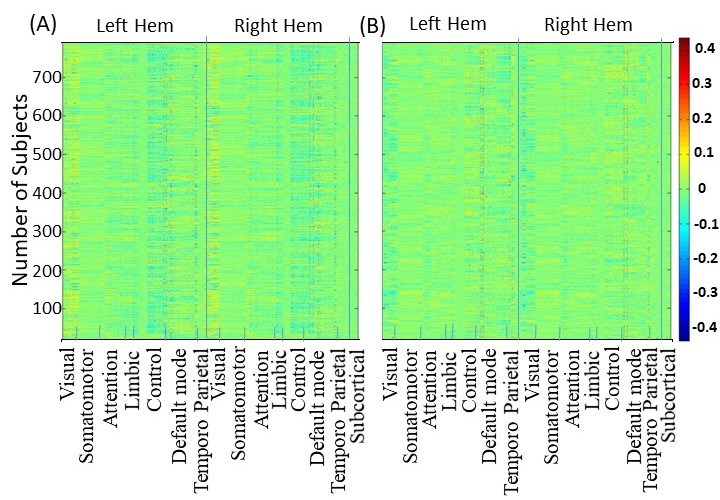

Figure S1. (A) Spatial distribution of weights of the second common COBE component of 788 subjects across the 7 networks (in both hemispheres) and 19 subcortical areas. (B) Same as (A) for the third common COBE component. Inter-individual variation in the distribution of weights is higher compared to the first common COBE component (figure 2C).

*Determination of Optimal Threshold*

We also tested different thresholds for selecting the subset of subjects whose pattern were dissimilar in the common correlation matrix. The thresholds were 0.65, 0.75 (as in MCD), 0.80, 0.85, and .90. We repeated the procedure to obtain the weights averaged across the subjects for the 17 cortical networks and 19 subcortical areas (refer methods section) for all the threshold groups of subjects. The number of subjects in the subset of each threshold was 55, 107 (as in MCD), 280, 423, and 689. Threshold ≤ 0.60 were not considered because that included only 13 subjects. On average, 92% of the total number of subjects included in a given threshold were also included in the higher thresholds. Figure S2 illustrates the weights of each group corresponding to a threshold. It is clear that the subset of subjects obtained with a threshold of 0.65 has similar weights as MCD (obtained with a threshold of 0.75) across all the networks. The weights with a threshold of .70 are similar to .75 and are therefore not shown in Figure S2. Similarly, the subset of subjects with threshold more than .75 also had similar weights like the set of subjects found with a threshold of .90. This led us to opt 0.75 as the optimal threshold and consider subjects in MCD as our study group.

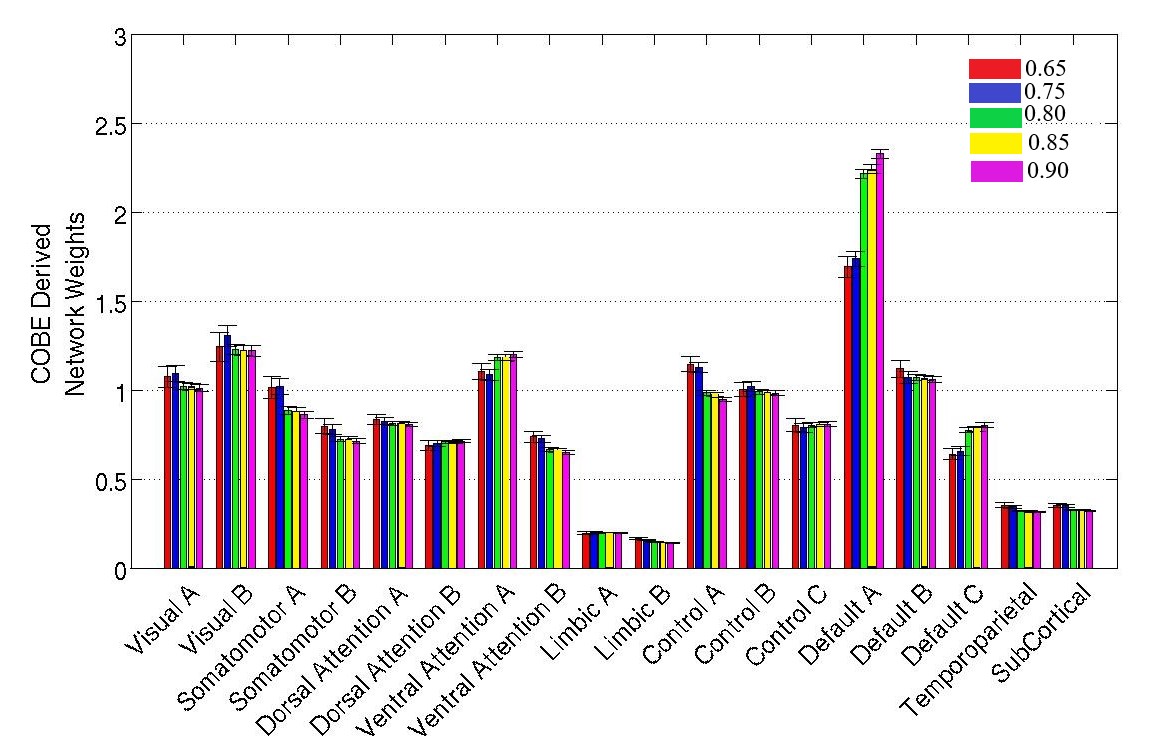

Figure S2. Distribution of weights across the 17resting state networks and subcortical areas for a subset of subjects with different threshold values (shown in different colors). Weight distribution for threshold 0.65 is similar to 0.75 and similarly, weight distribution for thresholds above 0.75 are also similar. So, the threshold of 0.75 seems to be an optimal choice.

From another perspective, we would like to mention that the subgroups that were formed for thresholds below 0.75 (see supplement) manifested similar differences in the scores of behavioral measures. For example, 55 subjects formed the group when the threshold was 0.65. The subjects in this group also reflected a significantly higher consumption of marijuana, illicit drugs, alcohol and tobacco and a predisposition towards problems in antisocial personality. Later we found these subjects to be a part of MCD (optimal threshold of 0.75), and therefore the behavioral results for thresholds below .75 were not highlighted. Similarly, a subset of subjects with higher thresholds (> 0.75) had no significant differences in the scores of behavioral measures. For example, 423 subjects formed the subset when the threshold was 0.85 and eventually the differences in the scores of behavioral measures did not pass the significance level.

As an alternative check we also wanted to see if correlations based on the COBE components were better than the correlation based on the average of the connectomes in separating the groups. For this, we averaged the functional connectivity of four runs of a subject’s fMRI to form the average connectome per subject. We then correlated the lower triangular matrix of every subject’s connectome to form the 788 x 788 averaged-correlation matrix (similar to common-correlation matrix). Next, we repeated our analysis as described in methods section. Since it was difficult to justify any optimal threshold, we adopted two thresholds (0.75 and 0.85) from our COBE analysis to form the MCD groups. Comparison of behavioural measures across the two groups (MCD and CS) for both thresholds did not fetch any significant differences. This could be expected as feature extraction and selection are key factors in model reduction, classification and pattern recognition problems. Thus, the COBE pipeline was helpful in pointing out meaningful differences in rs-fMRI features within subgroups of healthy individuals that had the distinctions translated to behavioral measures as well.

*Behavioral Measures across two groups*

***Table S1****.* Averaged scores of 68 HCP behavioral measures for the MCD group (n = 107) and the control group (n = 681). The significant behavioral measures are bolded.

| S.No | Behavior Category | Behavior  Name | HCP  Field Name | Mean ± Standard error | |
| --- | --- | --- | --- | --- | --- |
|  |  |  |  | MCD  (n = 107) | CC  (n =681) |
| 1 | Cognition | Visual memory | PicSeq_Unadj | 111 ± 9.73 | 112 ± 10.46 |
| 2 |  | Cognitive flexibility | CardSort_Unadj | 17.57 ± 4.23 | 17.27 ± 4.68 |
| 3 |  | Flanker Test | Flanker_Unadj | 111 ± 13.67 | 112 ± 13.16 |
| 4 |  | Fluid intelligence | PMAT24_A_CR | 117 ± 12.03 | 116 ± 10.33 |
| 5 |  | Reading | ReadEng_Unadj | 116 ± 9.46 | 117 ± 9.23 |
| 6 |  | Vocabulary | PicVocab_Unadj | 119 ± 10.35 | 117 ± 10.20 |
| 7 |  | Processing Speed | ProcSpeed_Unadj | 111 ± 10.62 | 111 ± 11.15 |
| 8 |  | Delay discounting | DDisc_AUC_40K | 0.96 ± .03 | 0.95 ± .03 |
| 9 |  | Spatial orientation | VSPLOT_TC | 35.35 ± 2.98 | 35.82 ± 2.77 |
| 10 |  | Attention Sensitivity | SCPT_SEN | 0.47 ±.28 | 0.52 ±.28 |
| 11 |  | Attention Specificity | SCPT_SPEC | 2.33 ± 3.46 | 2.78 ± 3.73 |
| 12 |  | Episodic Memory | IWRD_TOT | 16.05 ± 4.17 | 15.25 ± 4.33 |
| 13 |  | Working Memory | ListSort_Unadj | 114 ± 15.60 | 115 ± 15.27 |
| 14 | Alertness | Cognitive Status | MMSE_Score | 4.67 ± 2.88 | 4.58 ± 2.65 |
| 15 |  | Sleep quality (PSQI) | PSQI_Score | 28.91 ± 1.02 | 29.03 ± .98 |
| 16 | Sensory | Odor identification | Odor_Unadj | 1.80 ± 0.06 | 1.80 ± 0.06 |
| 17 |  | Pain Interference Survey | PainInterf_Tscore | 96 ± 14.06 | 94 ± 14.07 |
| 18 |  | Taste intensity | Taste_Unadj | 45.30 ± 7.21 | 45.20 ± 7.24 |
| 19 |  | Contrast Sensitivity | Mars_Final | 110 ± 9.23 | 110 ± 9.09 |
| 20 | Subject Information | Age | Age_In_Years | 27 ± 3.46 | 28 ± 3.72 |
| 21 |  | Gender  (*r =*$\frac{Number of Male}{Number of Female}$) | Gender | ***r =* 2.24** | ***r =* 0.90** |
| 22 | Health  and Family History | Height | Height | 76 ± 11.38 | 75 ± 10.23 |
| 23 |  | Weight | Weight | 126 ± 13.73 | 122 ±13.10 |
| 24 |  | Body Mass Index | BMI | 25.50 ± 4.38 | 25.16 ± 3.97 |
| 25 |  | Systolic Blood Pressure | BPSystolic | 173 ± 35.16 | 164 ± 33.88 |
| 26 |  | Diastolic Blood Pressure | BPDiastolic | 69.03 ± 3.88 | 67.58 ± 3.86 |
| 27 | Emotion | Agreeablenes | NEOFAC_A | 30.99 ± 6.65 | 30.85 ± 5.95 |
| 28 |  | Openness | NEOFAC_O | 15.97 ± 7.02 | 16.25 ± 7.30 |
| 29 |  | Conscientiousness | NEOFAC_C | 33.42 ± 6.02 | 34.61 ± 5.75 |
| 30 |  | Neuroticism | NEOFAC_N | 28.75 ± 6.34 | 28.58 ± 6.17 |
| 31 |  | Extraversion | NEOFAC_E | 31.69 ± 4.96 | 32.23 ± 4.74 |
| 32 |  | Emotion Recognition – Total | ER40_CR | 6.81 ± 1.16 | 6.85 ± 1.15 |
| 33 |  | Emotion Recognition –  Angry | ER40_ANG | 7. 16 ± 1.32 | 7. 19 ± 1.17 |
| 34 |  | Emotion Recognition –  Fear | ER40_FEAR | 7.95 ± .25 | 7.96 ± .20 |
| 35 |  | Emotion Recognition –  Happy | ER40_HAP | 6.84 ± 1.16 | 6.93 ± 1.13 |
| 36 |  | Emotion Recognition –  Neutral | ER40_NOE | 6.74 ± 1.07 | 6.80 ± .97 |
| 37 |  | Emotion Recognition –  Sad | ER40_SAD | 35.51 ± 2.77 | 35.71 ± 2.44 |
| 38 |  | Anger –Affect | AngAffect_Unadj | 46.33 ± 7.97 | 45.94 ± 7.73 |
| 39 |  | Anger – Hostility | AngHostil_Unadj | 52.07 ± 7.25 | 51.36 ± 7.91 |
| 40 |  | Anger – Aggression | AngAggr_Unadj | 50.01 ± 6.96 | 49.87 ± 7.83 |
| 41 |  | Fear – Affect | FearAffect_Unadj | 54.80 ± 9.45 | 51.37 ± 8.51 |
| 42 |  | Fear – Somatic Arousal | FearSomat_Unadj | 50.20 ± 8.35 | 49.64 ± 8.47 |
| 43 |  | Sadness | Sadness_Unadj | 48.04 ± 8.44 | 47.35 ± 7.94 |
| 44 |  | Life Satisfaction | LifeSatisf_Unadj | 50.69 ± 8.15 | 50.28 ± 7.82 |
| 45 |  | Meaning & Purpose | MeanPurp_Unadj | 51.89 ± 8.42 | 52.01 ± 8.79 |
| 46 |  | Positive Affect | PosAffect_Unadj | 54.70 ± 9.35 | 55.15 ± 9.19 |
| 47 |  | Friendship | Friendship_Unadj | 48.48 ± 8.82 | 48.32 ± 8.86 |
| 48 |  | Loneliness | Loneliness_Unadj | 51.67 ± 9.71 | 51.60 ± 9.40 |
| 49 |  | Perceived Hostility | PercHostil_Unadj | 48.82 ± 7.70 | 48.07 ± 8.48 |
| 50 |  | Perceived Rejection | PercReject_Unadj | 48.46± 7.86 | 48.15± 8.36 |
| 51 |  | Emotional Support | EmotSupp_Unadj | 51.01 ± 8.42 | 50.86 ± 8.13 |
| 52 |  | Instrument Support | InstruSupp_Unadj | 51.88 ± 8.50 | 50.87 ± 8.69 |
| 53 | Psychiatric and Life function | Aggressive | ASR_Aggr_Pct | 3.37 ± 2.56 | 3.11 ± 2.37 |
| 54 |  | Anxious/Depressed | ASR_Anxd_Pct | 54 ± 6.03 | 53.62 ± 5.51 |
| 55 |  | Attention Problems | ASR_Attn_Pct | 54 ± 6.23 | 52 ± 4.40 |
| 56 |  | Critical Items | ASR_Crit_Raw | 2.73 ± 2.24 | 2.31 ± 2.08 |
| 57 |  | Externalizing | ASR_Extn_T | 48 ± 10.63 | 47 ± 10.39 |
| 58 |  | Internalizing | ASR_Intn_T | 52.78 ± 8.77 | 48.18 ± 8.45 |
| 59 |  | Intrusive | ASR_Intr_Pct | 12.08 ± 8.01 | 8.23 ± 6.19 |
| 60 |  | Thought, attention and other problems | ASR_Tao_Sum | 6.90 ± 4.50 | 6.18 ± 4.26 |
| 61 |  | Antisocial Personality problems | DSM_Antis_Pct | **53.60 ± 12.5** | **52.18 ± 5.39** |
| 62 |  | Anxiety problems | DSM_Anxi_Pct | 5.50 ± 4.91 | 5.61 ± 5.21 |
| 63 |  | Depressive problems | DSM_Depr_Pct | 53 ± 5.59 | 53 ± 6.03 |
| 64 |  | Hyperactivity problems | DSM_Hype_Raw | 4.16 ± 3.64 | 3.51 ± 3.08 |
| 65 |  | Inattention problems | DSM_Inat_Raw | 53 ± 4.89 | 52 ± 3.64 |
| 66 | Substance Use | Alcohol use - 7 days | Total_Drinks_  7_Days | **10.11 ± 1.00** | **4.84 ± 0.26** |
| 67 |  | Tobacco use - 7 days | Total_Any_  Tobacco_7_Days | **18.05 ± 3.26** | **5.97 ± 0.76** |
| 68 |  | Times used Marijuana | SSAGA_Mj_Times_Used | **4.59 ± 0.04** | **1.35 ± 0.06** |
| 69 |  | Times used - cocaine, hallucinogen, opiates, sedatives, stimulants) | SSAGA_Times_  Used_Illicits | **2.19 ± 0.16** | **0.53 ± 0.04** |

*Correlation between behaviors*

The discussion notes that previous studies have reported correlations between substance use and problems in antisocial personality. Consequently, a natural question arises about the existence of any strong correlation between problems in antisocial personality and the other 4 behaviors (Marijuana intake, Illicit drug use, alcohol use, and tobacco use) for the 3 sample data (All subjects, CC and MCD group). However, none of the behaviors had a strong correlation across all the three samples, though the correlation value between antisocial personality and illicit drug use was fairly better than other correlations (Figure S3)

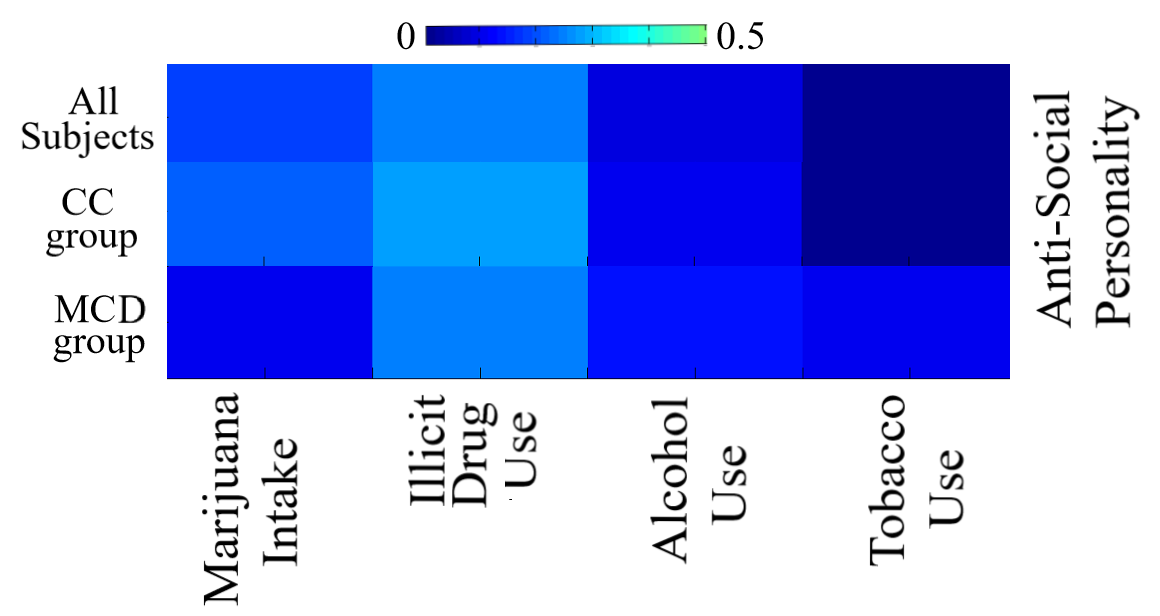

Figure S3. Correlation between the scores of problems in anti-social personality with the scores of the other four behaviors – Marijuana intake, Illicit drug use, Alcohol use, and Tobacco use. We tested this correlation for all subjects, CC and MCD group.

*Correlation of individual COBE variation with behaviors for MCD*

Since individuals with dissimilar COBE component shared common features to form the MCD subgroup, it would be interesting to explore whether the variation of the common COBE component within the subjects in the group (MCD) are associated to the five behavioral measures (Marijuana intake, Illicit Drug use, Alcohol and Tobacco use, and Antisocial personality problems). To do this, we obtained the variance of the common COBE component of each individual in the MCD group. We then evaluated the correlation of the obtained values of variance with the scores of the five behavioral measures (Figure S4). We found none of the behaviors to have a strong correlation with the variance, though antisocial personality problem faired better than the others. With that said, it remains difficult to comprehend why individuals with dissimilar COBEs themselves formed a coherent subgroup.

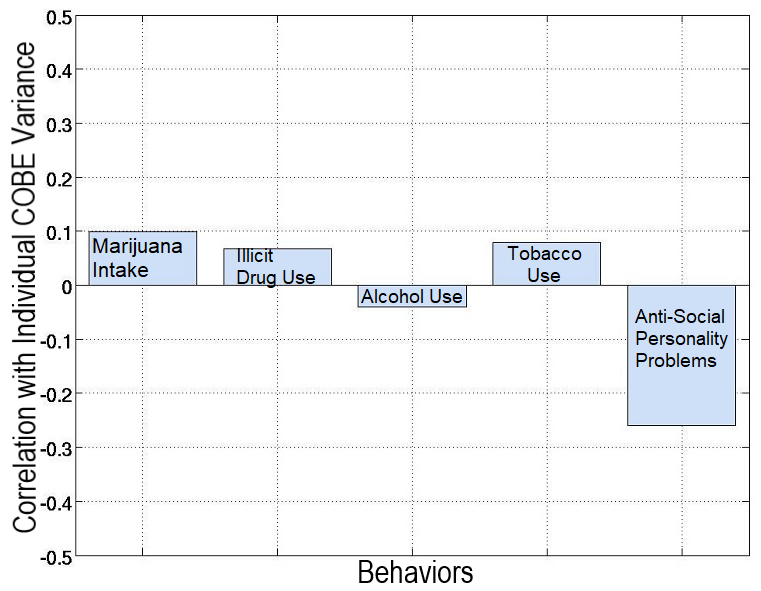

Figure S4. Correlation between variance of the common COBE component of each individual in the MCD group with the scores of the five behavioral measures.
